# Thalamic Influence Over Adaptive Cortical Dynamics Across Conscious States

**DOI:** 10.1101/2024.07.02.601788

**Authors:** Eli J. Müller, Brandon R. Munn, Giulia Baracchini, Ben D. Fulcher, Vicente Medel, Michelle J. Redinbaugh, Yuri B. Saalmann, Bingni W. Brunton, Steven L. Brunton, James M. Shine

## Abstract

The human brain must support both stable and flexible neural dynamics in order to adapt to changing contexts that are inherently non-linear. The thalamus has been linked to the coordination of these opposing dynamical regimens in the cerebral cortex, however existing methodological approaches have not integrated sufficient neurobiological details with a sensitive measure of neural dynamics that permits sensitivity to time-series non-linearities. Inspired by the field of fluid dynamics, we use a novel approach to show that spontaneous fMRI data exhibits non-trivial fluctuations in predictability over time, akin to a river that has sections of smooth and predictable (laminar) versus rough and unpredictable (non-laminar) fluid flow. We use a combination of pharmacological fMRI, macaque electrophysiology and a large-scale biophysical model of the thalamocortical system to provide robust evidence that the thalamus provides versatile control over globally linear dynamics in the cerebral cortex that characterize conscious states.

## Introduction

The human brain supports a wide-ranging behavioural repertoire, with dynamic, context-dependant recruitment of specialized neuronal assemblies emerging from coordinated interactions amongst tens of billions of cells. The computational capacities that support these behaviours are in large part related to this cellular complexity, however, this high-dimensional neural activity is of minimal adaptive benefit unless the neural activity that unfolds over time is both stable and yet flexible. In other words, neural dynamics must be reliable for repeated precise execution, but also rapidly adaptable to changing contingencies. How the intricate organization of the brain supports these opposing capacities remains to be understood.

The thalamus is a highly-conserved subcortical structure that has been argued to play a central role in shaping and constraining the flexible neural dynamics that characterise the conscious, awake brain^1–3^. At the microscale, there is evidence that specific cell types in the thalamus act to relay information from sensory systems^4,5^, coordinate motor actions^6^, and also to integrate and distribute information from subcortical structures^7–9^ to decentralized corticothalamic networks. At the macroscale, recent work has shown that diffusely-projecting subtypes of thalamocortical nuclei promote variability of dynamics across the cerebral cortex^7,8,10^ through the instantiation of quasi-critical brain state regimes^11^ which support shifts in cognition^12^ and consciousness^13–16^. These emerging perspectives place the thalamus in a unique topological position within the brain, wherein it is able to both drive the cerebral cortex with targeted inputs, but also to modulate on-going cortical dynamics through diffuse projections^7^. Despite this privileged position, to date there are few direct empirical links between the thalamocortical system and precise dynamic trade-offs characteristic of adaptive systems^1^.

A primary reason for this lack of direct evidence is that few existing neuroimaging analysis techniques are capable of providing a time-resolved estimate of the stability (or flexibility) of brain state dynamics that is sensitive to the inherently non-linear waxing and waning nature of our waking lives^17^. Most of the current statistical approaches have either been designed to capture distributional moments of neural time-series or have heavily relied on assumptions of stationarity^18,19–25^. While some approaches have softened assumptions of stationarity – e.g., by carving data into shorter windows that are swept over time^26,27^ – these techniques are typically model-based, which makes the detection of non-linear shifts in brain states inherently difficult. Other approaches have used inventive means for tracking functional neuroimaging data at a fine temporal resolution^28–30^, however the direct interpretation of these measures with respect to stability and flexibility (which naturally require longer epochs of data to quantify) is non-trivial.

Fortunately, there are existing methods from the field of fluid dynamics that are well-suited to this class of problems, however they require a subtle change in perspective. By way of analogy, if we consider brain dynamics as a flowing river, we need to move from tracking the topography of the river – which is admittedly still highly informative – to characterising the flow of water at different sections of the river: one section might be calm and largely laminar, whereas another might be full of turbulent swirling eddies and so non-laminar (Fig. 1a). Importantly, these local dynamics provide fundamentally different constraints over the local environment, permitting smooth (stable) or disturbed (flexible) flow of local objects, respectively. This dynamics-focussed perspective has led to key insights in fields such as fluid mechanics, mechanical engineering, economics, and sociology^31–33^, and here we advocate for its application to problems in neuroscience.

**Figure 1.**
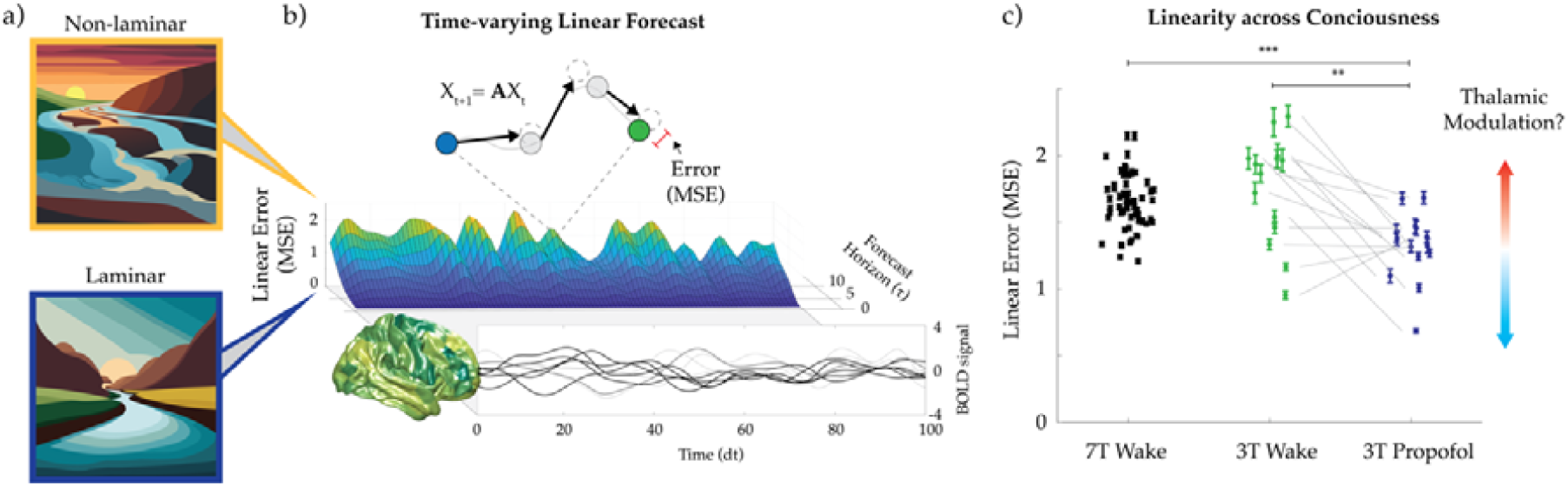
Linear dynamics in large-scale brain recordings. a) Depiction of laminar and non-laminar river flows forming an analogous axis for large-scale brain dynamics – this analogy provides a useful means for appreciating the approach used in our experiment. b) Top: Schematic showing how a global linear model can be compared to ground-truth timeseries as a time-varying measure of linearity – here, a low-dimensional representation of changes in the BOLD signal across the cerebral cortex is used to predict changes to the state of the brain in the next time point (black lines to dotted circles); the actual brain state (closed circles) is recorded, and the distance between these two states in state space is quantified as the “Error” of the model – linear dynamics are associated with consistently low Error values, whereas non-linear dynamics have variable Error values. Bottom: A Horizon plot showing the time-varying linear forecast performance in 7T resting-state human BOLD^43^ from an example subject – note the large fluctuations in Linear Model Error (MSE), particularly at longer delays (Forecast Horizon,_1_). c) 7T human resting-state linear prediction performance (N=59; black), 3T human resting-state linear prediction performance (N=14; green), 3T human resting-state under propofol anesthesia linear prediction performance (N=14; dark blue).

There are several significant historical parallels between the fields of neuroscience and fluid dynamics in the development of measurement techniques that further motivate advances in data-driven modelling approaches. Both disciplines study complex systems that are high-dimensional, non-linear, and inherently multi-scale, both in space and time, yet both also exhibit dominant spatiotemporal patterns of activity that are meaningful for scientific understanding and engineering^31,34,35^. Importantly, experimental measurements in each field have followed a similar technological trajectory: recordings were first single point (hot-wire anemometry and sharp electrodes), followed by simple arrays (many hot-wires and electrode arrays), and more recently spatial and temporal fields resolved by imaging (laser-based particle image velocimetry PIV^36–38^ and magnetic-resonance-based functional brain imaging^39^). Accordingly, mathematical modelling techniques in fluids have evolved from quantifying statistics to extracting spatiotemporal dynamics^40,41^. We note that—with a delay of several decades—a similar progression is now occurring in neuroscience^42^. For this reason, there is a tremendous opportunity to leverage modern dynamical systems modelling techniques to gain insights into neural computation.

In this paper, we repurpose an approach from fluid dynamics for application to functional neuroimaging data. Specifically, we develop a measure of timeseries stability – akin to fluid laminarity – by tracking the capacity of a simple linear dynamic model to accurately predict successive time-points. We show that neural recordings transition through periods of increased linear predictability that irregularly fluctuate through periods of decreased linearity over time (Fig. 1b). The spatial and temporal structure of these linear dynamics reveals coherence maps and intrinsic timescales that are not captured by null models that either scramble temporal order or recreate timeseries based on the covariance and Fourier power spectrum of the original data. We then reveal how pharmacological manipulation of arousal, via propofol-induced anesthesia, results in heightened linearity in human neuroimaging data (Fig. 1c). Next, we demonstrate how targeted and diffuse thalamocortical subtypes can both promote or diminish linearity, respectively. We underscore the importance of thalamocortical interactions in a detailed biophysical model which further predicts matrix thalamic stimulation re-instantiates deviations from linear dynamics concomitant with conscious arousal, which we then confirm by applying our approach to multielectrode cortical recordings from an anesthetized macaque that was awoken by electrical stimulation of the diffuse thalamic projections within the central lateral thalamus^14,15^. In this way, we reveal the crucial role of the thalamus in shaping and constraining global linear activity patterns distributed across the cerebral cortex, supporting an essential feature of stable, and yet adaptive and flexible cognition.

## Results

### Linear dynamics in large-scale cortical recordings

To quantify the time-varying stability of brain state dynamics, we applied a simple linear model estimation technique (Fig. 1b; Supp. Fig. 1) to a high spatiotemporal resolution human 7T fMRI dataset captured during the resting state. First, a global linear propagator tracking moment-to-moment shifts in a recording is generated for each subject’s timeseries by utilizing the singular-value-decomposition (See Supp. Fig. 1a). Then, for every timepoint in the recording, the linear propagator is used to forecast state dynamics at a set of future timepoints (τ; 1-10 TR) – note that we explicitly chose to fit a global propagator (i.e., we did not redefine the propagator in a sliding-window fashion). The linear dynamics predicted by the global propagator are then compared to the ground-truth dynamics via the mean-squared-error, providing a time-resolved read-out of how effective (or not) the linear model was at predicting the upcoming brain state. We denote this term as the Forecast Horizon of the linear model prediction (Fig. 1b) and the resultant graph as a Horizon plot.

We reasoned that, if the resting brain were stable (i.e., linear) throughout the recording, then we should observe relatively strong, consistent predictability throughout the entire recording session. In stark contrast, the Horizon plot in Fig. 1b shows that neural dynamics cycle through periods of increased and decreased linearity over the course of a recording session. This observation is at direct odds with the typical assumption of stationarity that is applied in standard neuroimaging analyses^21^. The results are observed for every individual in the dataset (N = 59; Supp. Fig. 2), robust across validation datasets (which had different temporal resolutions; see Supp. Fig 2) and do not correlate with frame-wise displacement (r = 0.01 +−0.03 across 10τ). Crucially, these results extend previous findings of strong linear dynamics in human neuroimaging^44–46^ and large-scale electrophysiological recordings^46,47^, by showing substantial yet irregular deviations from linear predictability in conscious states that is pervasive across subjects.

### Adaptive timescales and dynamic brain modes

Intrinsic timescales of the brain have been of interest since the discovery of oscillatory dynamics in the first human EEG recordings^48^. Human neuroimaging studies have revealed a hierarchy of temporal and spatial autocorrelation scales across the cortex^49^ and that these capture a large number of existing topological metrics^22^. Linear flow analysis enriches this approach by quantifying intrinsic modal timescales that drive linear dynamics in neural activity – and which may be recruited in an adaptive and time-varying manner reflecting changing cognitive demands. To do this, we leveraged another method commonly used in the field of fluid dynamics – *Dynamic Mode Decomposition* (DMD) – that is designed to efficiently characterize the dynamics of time-varying processes through eigendecomposition of the linear propagator matrix^31,34,35,50^, reformulating the linear dynamics as a function of spatiotemporal modes ^32,35,50^. Each mode contains four key characteristic properties: the spatial eigenvector defines a spatial pattern of coherent activity, and a corresponding spatial pattern of delays relative to each of these coherence patterns – called a dephasing map. In addition, each mode has a corresponding eigenvalue defining its oscillatory frequency, and an exponential gain parameter – which determines whether a mode will grow or decay in time (i.e., captures its temporal stability) – representing two key dimensions of the system’s timescales.

By applying DMD to each subject’s fMRI recording, we formulate a subject-specific set of spatiotemporal modes capturing on-going linear dynamics. In order to gain insight into how these dynamic modes relate across all subjects, we leverage a two-pronged clustering approach to aggregate modes across timescales, and then spatial coherence, respectively. First, we consider timescales by utilizing *k*-means clustering of the eigenspectrum collated across all subject recordings. A peak in adjusted mutual information between clustering repetitions was then used to select *k* = 5 clusters (1000 repetitions; see Supp. Fig. 4), resulting in 5 distinct timescale groupings – the corresponding average of the coherence maps within these clusters (Supp. Fig. 4). Timescale cluster 1 shows somatomotor and visual activation antiphase with frontal, parietal and temporal cortices across a broad range of frequencies – spanning much of the time resolution of BOLD recordings. These modes are also weakly damped – i.e., have a strong gain relative to the other temporal clusters. Timescale cluster 2 shows visual, premotor, and dorsal lateral prefrontal antiphase with other cortices at slower frequencies (< 0.03 Hz; Fig. 4 g-h middle) and with a stronger damping rate, i.e., weaker gain, than cluster 1. And timescale cluster 3 shows strongly damped temporal cortex activity at frequencies < 0.02 Hz.

Next, we considered the spatial patterns of coherence, defined by each mode, aggregated across subjects and recording sessions to define consistent modal groupings^50^. Again, we leveraged *k*-means clustering (adjusted mutual information peaked between clustering repetitions for *k* = 5; See Supp. Fig. 4): Figure 2m shows 3 cluster centroids and their defining coherence patterns – i.e., the spatial patterns of the linear modes across all subjects fall within distinct classes. These classes show a significant overlap with resting-state networks used broadly in the literature^51^ – including the default mode, and somatomotor and visual networks. We further find that the damping rates and frequencies of the modes within each class have comparable probability distributions – demonstrating these spatial classes did not demonstrate unique linear timescales at this granularity in resting-state.

**Figure 2.**
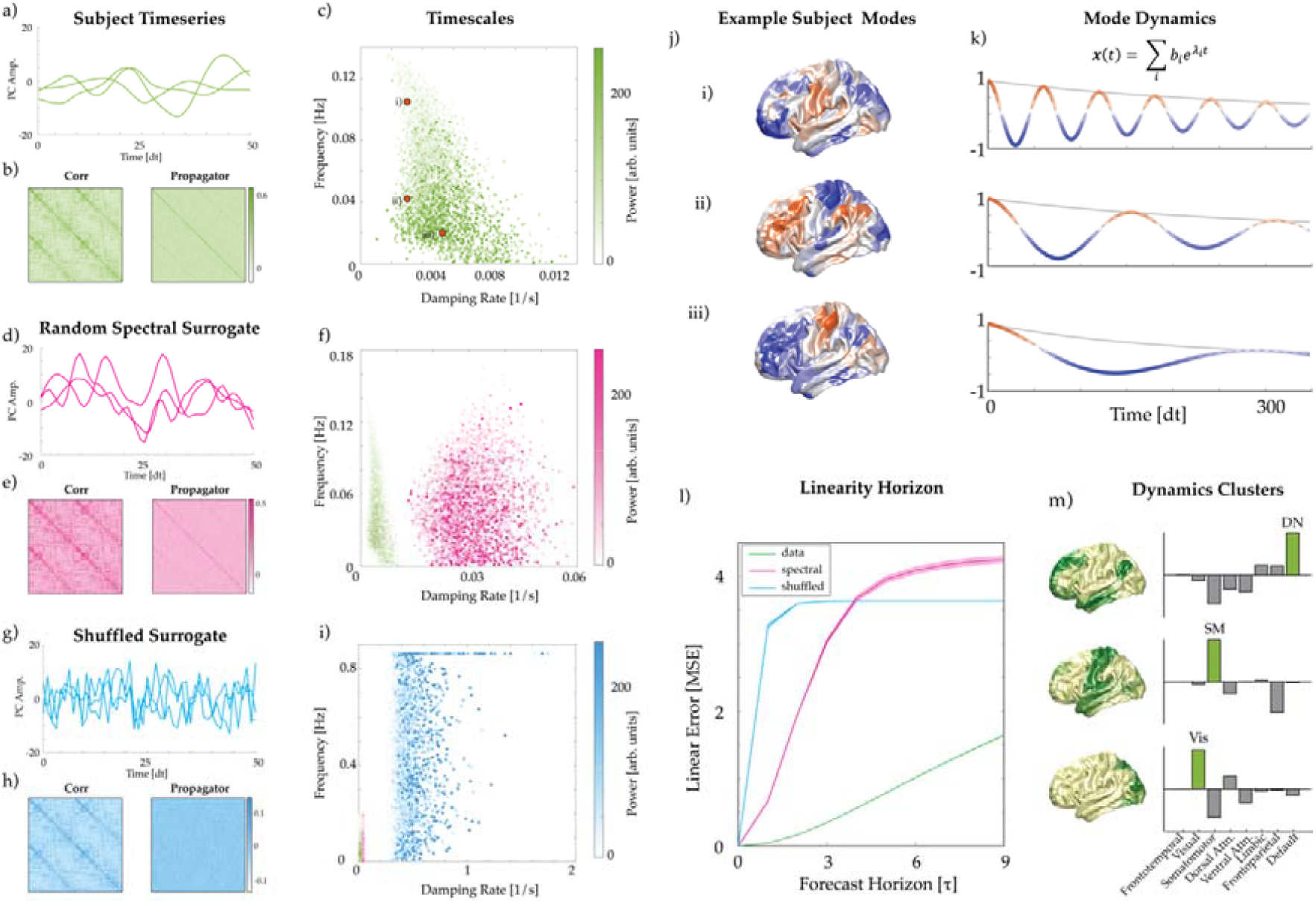
Spatiotemporal modes of global linearity. a) Example subject time-series of first 3 principal components. b) Example subject correlation matrix (left) and linear propagator matrix (right) capturing linear dynamics. c) Timescale properties of dynamic modes defined by the linear propagator of 7T resting-state data (N=59). Colour bar shows the total power within each mode across the entirety of the recording. Orange dots correspond to the example subject’s dynamic modes in j). Random Spectral Surrogate is generated from a white-noise timeseries with a covariance and Fourier power spectrum matching the empirical data in a-b). d) Principal component time-series for the random spectral surrogate. e-left) Random spectral surrogate correlation matrix, (e-right) linear propagator matrix, and (f) dynamic mode timescales. Shuffled Surrogate is generated by uniformly randomizing each timepoint of the regional timeseries – i.e., only randomizing time and not space. g) Principal component time-series for the shuffled surrogate. h) Left: shuffled surrogate correlation matrix; Right: linear propagator matrix, and (i) dynamics mode timescales. j) Example subject dynamic modes capturing the most (j-i), median (j-ii), and least (j-iii) amount of globally-linear dynamics and (k) these modes’ corresponding temporal dynamics. l) Average linear errors as a function of future time points across all (N=59) subjects (green), and all corresponding Random Spectral (pink), and Shuffled (blue) surrogates (p < 0.001; line = mean across subjects; shaded error bar = variance across subjects). m) k-means spatial coherence clusters of the dynamic modes across all subjects (N=59) - showing 3 clusters (m-left) of k=5 – see Supp 3 for all clusters. (m-right) Cluster power within each of the 7 resting-state networks^51^.

### Empirical fluctuations are distinct from surrogate models

To further investigate the fluctuations in linear dynamics observed in resting state fMRI data, we contrast these against two surrogate timeseries that conserve distinct features of the empirical fMRI data. The first, referred to as the *spectral surrogate*, consists of white-noise timeseries imbued with a covariance and Fourier frequency spectrum matching those of the empirical data (i.e., a stationary surrogate^21^; see Supp. Fig. 1d). Crucially, this surrogate shares the empirical correlative structures between brain regions which forms the basis of many approaches in the neuroimaging literature^18^. In addition, this surrogate includes timescale information captured by the Fourier spectrum, which is predictive of future dynamics, since the sinusoidal modes comprising this time averaged measure oscillate across time. The second surrogate, referred to as the *shuffled surrogate*, is simply the initial empirical fMRI data uniformly scrambled in time – i.e., the regional activations relative to other regions at a given moment are conserved, matching empirical regional correlations, but *when* they occur with respect to any other activations in time is scrambled. Since the empirical data and both surrogates have the same correlation structure, they also share the same principal components (i.e., covariance defining regional activation patterns). Despite these similarities, Fig. 2b & 2c show that the temporal expression of these patterns no longer matches that seen in the empirical data (Fig. 2a).

Again, leveraging *Dynamic Mode Decomposition*, we find the empirical fMRI data are clearly distinct from the surrogates in two key ways: firstly, the fMRI data consist of slowly fluctuating, sustained, spatiotemporal dynamic modes (Fig. 2c), as compared to the surrogates. Due to its matched oscillatory spectrum, the spectral surrogate contains a similar distribution of modal frequencies (0-0.14Hz) to the empirical data (Fig. 2f). However, these oscillations are strongly damped, as the spectral surrogate mostly excludes moment-to-moment information predictive of future states with only the time-averaged Fourier modes contributing to continuity of its temporal trajectories. In contrast, the shuffled surrogate displays a much wider range of high frequency (0-0.8Hz) strongly damped modes than both the empirical data and the spectral surrogate (Fig. 2i). This is due to the shuffled surrogate’s time-point scrambling favouring fast and short-lived shifts in moment-to-moment activity.

Secondly, the empirical fMRI data contains significantly more linear dynamics than either surrogate (Fig. 2l). That is to say, the empirical fMRI data contain information in their moment-to-moment dynamics that is predictive of future globally-linear trajectories. Since both surrogates share the same covariance as the empirical data, and the spectral surrogate additionally includes Fourier spectral structure, we find these time-averaged features are not sufficient to capture the observed linear dynamics seen in the neural recordings. This finding was consistent at the subject level for all subjects (N=59; Supp. Fig. 2) and cross-validated in a 3T resting-state imaging dataset (N=14; Supp. Fig. 2^52^). To ensure that this effect was not caused by fixed patterns enduring within the timeseries, we calculated the time-averaged mean-squared-displacement (MSD) for successive lags – comparable to temporal autocorrelation of the entire brain state – and found the linear errors are significantly smaller than those expected from general variability within the signal (Supp. Fig 3). In short, we have presented a simple technique for clearly demonstrating that, despite substantial fluctuations between linear and non-linear flow, resting state fMRI data remains distinctly *more* globally-linear than either the spectral or shuffled surrogate.

How does this perspective of linear dynamics extend traditional investigations into the resting brain? Further insight can be gathered by exploiting the dynamic modes corresponding to the propagator used to predict the linear (or non-linear) temporal epochs (Fig. 2l). Figure 2j shows an example of three dynamic modes from a single subject’s recording: the dynamic mode with the largest average amplitude across the time series (Fig.2j-i), median (Fig.2j-ii), and the smallest (Fig. 2j-iii). The spatial coherence maps for these modes are shown in Fig. 2j and have corresponding temporal dynamics defined by a complex exponential (Fig. 2k). These modes can be collated across all subjects via clustering approaches (See Methods) and show spatial coherence groupings that recapitulate common resting-state networks^51^ (Fig. 2m; Supp. Fig. 4), consistent with previous findings^53^, and temporal groupings defining cross-regional dynamics at distinct frequencies and temporal stabilities – i.e., damping rates (Supp. Fig. 4).

Together these results clearly demonstrate that fMRI data contains predictable linear dynamics that are not observed in stationary surrogates – that is, the dynamics of human fMRI contains information in its moment-to-moment fluctuations that is predictive of its future dynamics. Despite this predictability, resting-state neuroimaging data also irregularly cycles through periods of predictable and unpredictable flow (Fig. 1b).

### Thalamic axis of cortical linearity

Armed with this simple yet effective approach, we were now able to test our original hypothesis: namely, that axes of anatomical variation in the thalamus facilitate shifts between stable and flexible dynamical modes in the brain. The thalamus itself is comprised of a rich diversity of cell types which have unique projection profiles to both the cerebral cortex and other subcortical structures^54–56^. Although there is no single schema that effectively captures the diversity of thalamocortical interactions^9,57–60^, recent empirical studies in rodents have demonstrated the utility of considering a non-binary continuous spectrum of thalamic cells that differ in terms of the topography and targets of their axonal arbours^61^. At one end of this continuum sit the calbindin-rich matrix nuclei, which have thalamocortical projections spanning multiple cortical areas, typically targeting superficial layers of agranular association cortices (Fig. 3a). This organization contrasts with the parvalbumin-rich core nuclei, which typically target granular layers of the cerebral cortex, providing feed-forward input to cortical pyramidal neurons^8,9^ (Fig. 3a-b).

**Figure 3.**
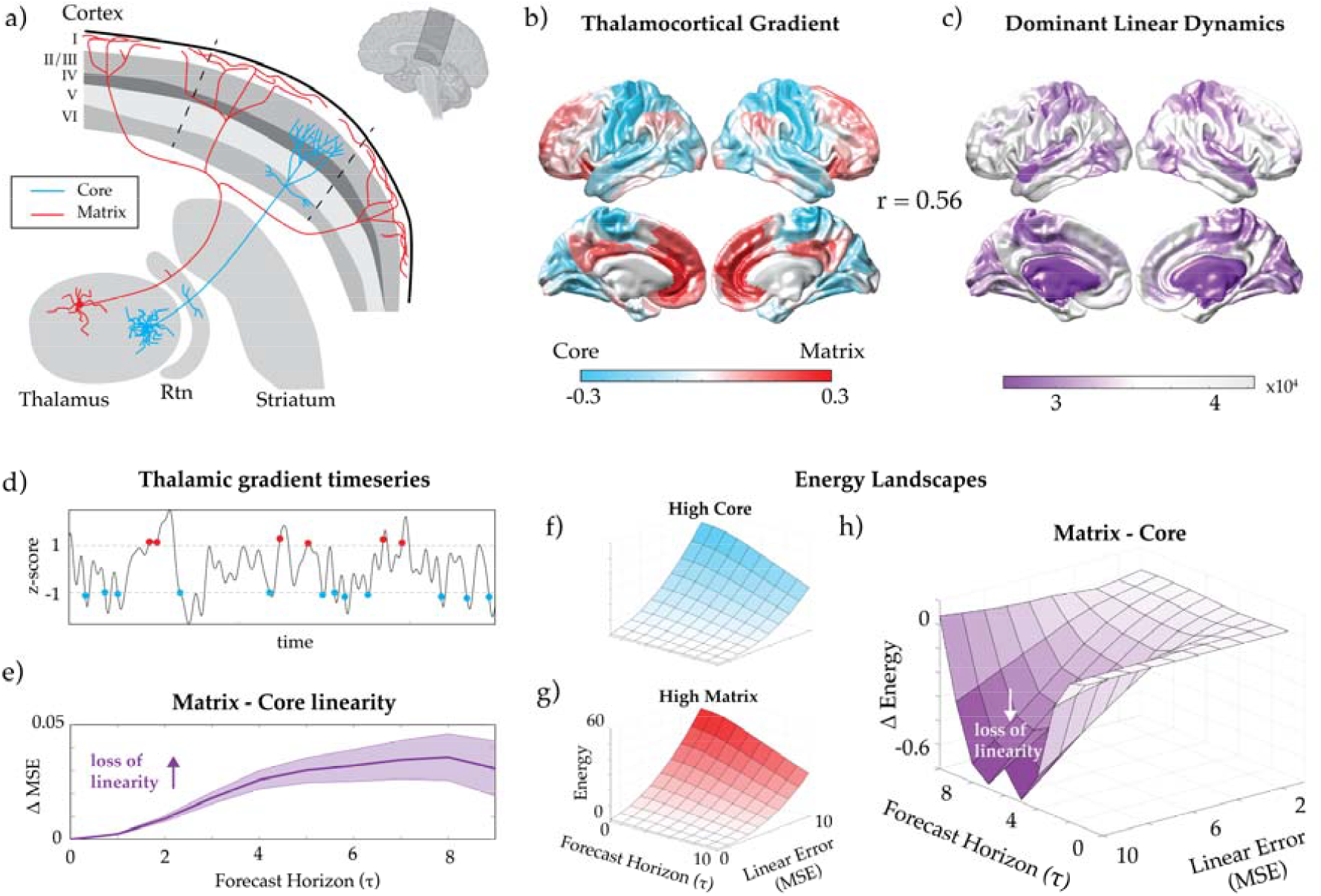
Thalamic axis of cortical linearity. a) Schematic of characteristic projection profiles of core and matrix thalamocortical projection profiles (adapted from ^54^). b) Core-Matrix thalamocortical expression gradient (adapted from ^8^). c) Population average cortical map showing regions of dominant linear dynamics 10 time-points into the future. d) Thalamic timeseries for each subject are projected along the core-matrix gradient and epochs showing one standard deviation above the mean (high matrix) and below the mean (high core) are collated. e) The difference of the mean-squared-error of the linear model following high matrix compared to high core thalamic activity. These mean-squared errors can be reformulated into corresponding energies^62^ (logarithm of inverse state probability) following high core (f) and high matrix (g) thalamic activity. h) The subtraction of the core energy landscape from the matrix energy landscape. Negative values indicate strong linearity following core activity as compared to matrix activity.

Although there is evidence of individual thalamic cells that contain axonal features of both core and matrix types (even in the same axon^54^), there are good reasons to consider a spectrum that extends between the two extreme phenotypes^3,8^. For one, simplifying assumptions at the microscale are required in order to conduct tractable work at the macroscale of neuroimaging^1,8,11,13^. Secondly, there is now robust evidence that core and matrix cell types also differ in their functionality^3^: recent empirical and theoretical work has shown that electrical stimulation of the central lateral thalamus – which itself is abundant with matrix nuclei – but not the ventral lateral thalamus (which is a predominantly core-rich nucleus) can drive recovery of consciousness in propofol-anesthetized macaque monkeys by leveraging the non-specific diffuse projection profiles of these matrix cells^11,13–16^. From these empirical vantage points, we thus ask: what type of cortical dynamics are facilitated by the thalamus, and how do these dynamics relate to the expression of core and matrix thalamic cell subtypes (Fig. 3)?

To answer this question, we applied a multitiered approach. First, we utilized the difference between the ground-truth data and the predicted linear dynamics – calculated for each region, each timepoint and each subject. We then sum these errors across regions to create a forecast horizon τ= 10 (ensuring sufficient time for autocorrelation to decay) across all time points and all subjects. This defines a cortical map revealing a spatial pattern of dominant globally-linear dynamics (Fig. 3c). Figure 3c shows that predictable linear flow was predominant in sensory cortices, and that these dynamics were strongly negatively correlated with the core-matrix expression gradient^8^ (r = −0.56; p < 10^−10^; Supp. Fig. 6a). That is, cortical brain regions with strong core thalamocortical projections also showed strong linear dynamics. However, both of these spatial maps also covary with the first principal component of the imaging data, which itself captures a significant proportion of the empirical timeseries’ overall covariance, thus making strong inferences about the thalamic involvement difficult.

To test our hypothesis more precisely, we next asked whether the thalamus was temporally related to fluctuations in the predictability of cortical dynamics. Specifically, a time-resolved measure of matrix thalamus (relative to core thalamus) was generated by projecting each subject’s thalamic BOLD timeseries onto a core-matrix expression gradient approximated from Allen Human Brain genetic expression data that corresponds to both extreme thalamic cell phenotypes (see methods; Fig. 3d)^8^. In this data, a positive weighting corresponds to high matrix activity relative to core regions of the thalamus (and *vice versa*). Next, for each subject, we identified peaks in this matrix-core timeseries that were one standard deviation above or below the mean (for high matrix and high core activity, respectively). Temporally averaging the prediction errors across all subjects and subtracting high core epochs from high matrix epochs clearly shows that increased linearity of cortical dynamics following peaks in core thalamic activity (Fig. 2e) – i.e., core thalamic activity precedes stable linear flow in the cortex.

To investigate these temporal fluctuations further, we leveraged a recent technique motivated by approaches in statistical thermodynamics that quantifies the likelihood of moment-to-moment state changes in a given timeseries by representing fluctuations as thermodynamic energy^62^. The likelihood for changes in linear flow (MSE) in the data is generated at each proceeding timepoint (forecast horizon, τ) and produces a corresponding energy landscape for the two thalamic conditions (Fig. 2f-g; see Methods for details). This approach strengthens our previous findings, in that strong core thalamic activity (relative to matrix thalamic activity) is followed by increased predictable dynamics (negative drop seen in Fig 2h) in the cerebral cortex (with a maximum ~3 seconds post-peak).

These results are in keeping with the targeted feed-forward projections from the core thalamus acting as a feed-forward propagator cortical activity, and the more diffusely-projecting axons from matrix thalamic nuclei acting more akin to a modulator of ongoing cortical activity^14–16,63^. Furthermore, previous work^8^ has shown that this anatomical axis also covaries with temporal variability of corticocortical connectivity, as captured by dynamic correlations in fMRI^8^. That is, cortical regions receiving more projections from the matrix thalamus showed a greater flexibility in their coordination with other cortical regions. Together, these findings demonstrate that the thalamus can provide both the stabilization of on-going cortical dynamics necessary for robust behaviours, but crucially, also mediate the flexibility of these cortical dynamics, which is essential for adaptive cognition. However, to strengthen the evidence for the thalamic control over cortical linearity, we require approaches that causally influence brain states.

### Anesthesia promotes linear flow in the cerebral cortex

We first sought to determine whether alteration of conscious brain states impacted the linear predictability of cortical dynamics. Given the highly susceptible, flexible and context-dependent nature of consciousness^64–66^, the critical importance of the thalamus in controlling dynamic brain states^1,14,67^ (Fig. 3), and armed with a technique for quantifying this impact (Figs. 1 & 2), we reasoned that awake state should be associated with irregular deviations from linear flow, which would then be diminished during anesthesia. To test this hypothesis, we extended our approach to an existing human neuroimaging resting-state fMRI dataset of 14 healthy volunteers (4 women; mean age 24 years, SD = 5^52^) who underwent propofol-induced anesthesia. As predicted, we found that anesthesia strikingly disrupted deviations from linear predictability compared to the awake state, with the linear forecasts performing significantly better across all subjects following propofol administration (blue line in Fig. 4d). This population-level observation can be better understood when viewing the Horizon plot of a specific subject recording, where the wake dynamics show periods of increased linear flow (Fig. 4b), which are equivalent to those observed for the same subject under propofol anesthesia (Fig. 4c). In contrast, the waking data is punctuated by periods of sharp deviation from linear flow that are not observed during propofol anesthesia (Fig. 4b). This observation thus provides confirmatory evidence for our hypothesis linking deviations from linear predictability to the conscious, waking state.

**Figure 4.**
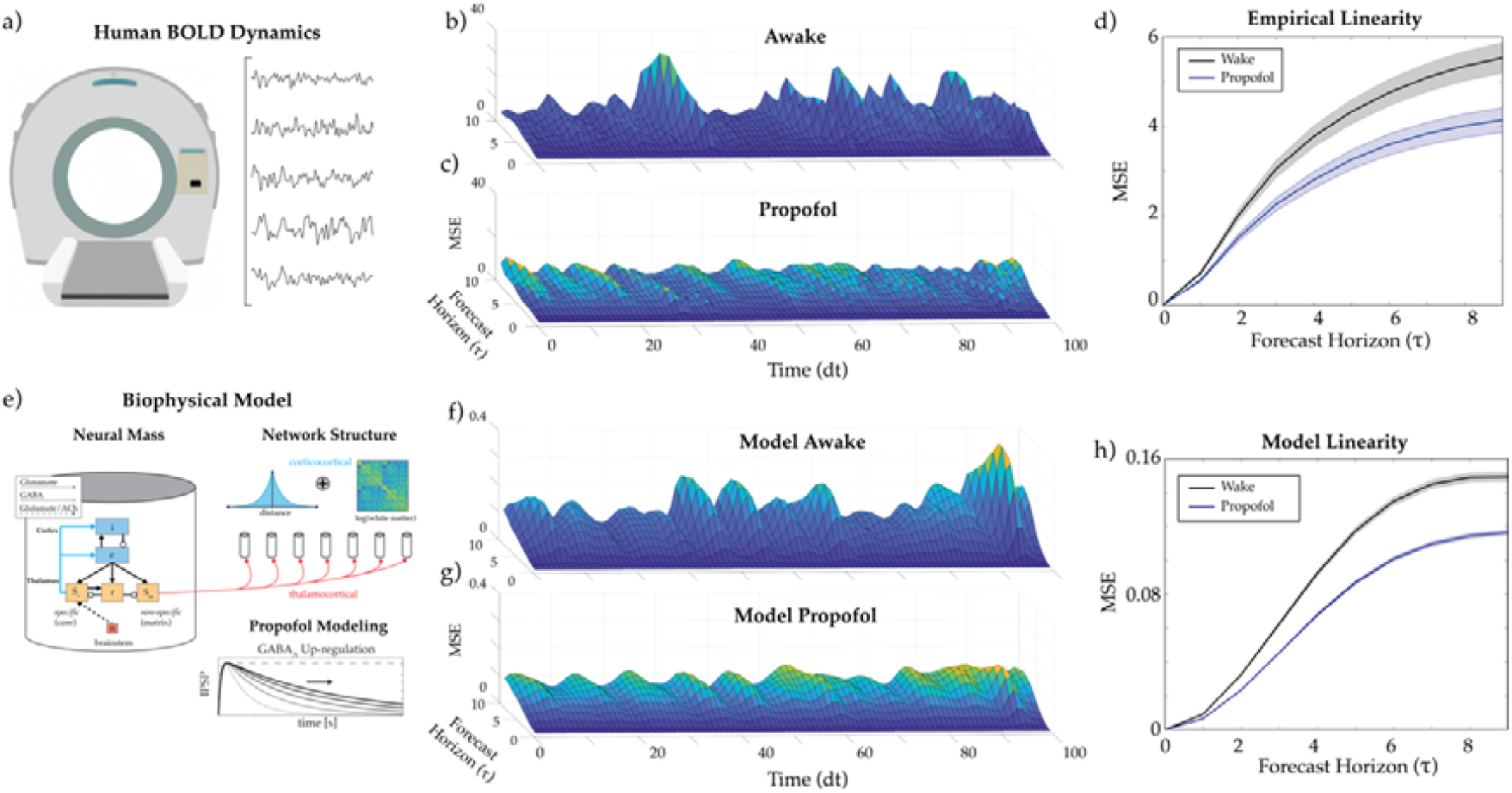
Linear flow across arousal. a). Schematic of fMRI timeseries. b-c) a Horizon plot of an example single subject resting-state human BOLD showing time-varying linear error during wake (b) and during propofol anesthesia (c). d) Population average linear error of BOLD resting-state data during wake (black) and propofol (dark blue) conditions averaged over each recording – shaded regions show standard-error-of-mean (note: this is a group-level summary of the data in Figure 1c). e) Schematic of large-scale biophysical neural mass model fit to human resting-state BOLD data^13^. As previous^13^, propofol is modelled as a prolongation of inhibitory-postsynaptic-potentials. f) Horizon plot showing time-varying linear error of the corticothalamic model timeseries - transformed using a haemodynamic response function to provide representative BOLD timeseries - fit to wake condition. g) Time-varying linear error of the corticothalamic model timeseries under the propofol condition. h) Average linear error of corticothalamic model during wake (black) and propofol (dark blue) conditions.

To further interrogate this empirical observation, we leveraged a state-of-the-art, large-scale biophysical neural mass model capable of describing consciousness modulation under propofol anesthesia, and its recovery following stimulation of the matrix thalamus^13^. This corticothalamic model includes realistic population-level neuroanatomy, non-linear neural responses, interpopulation connections, long-range white-matter connectivity, and dendritic, synaptic, cell-body, and axonal dynamics^11,13,68–73^. The biophysical constraints incorporated in this model allow putative physiological mechanisms to be explored which provide insights beyond those obtained via purely data-driven approach.

Applying our same approach to the wake and propofol conditions of this model’s timeseries – transformed using a haemodynamic response function to provide representative BOLD timeseries – we replicated our empirical observations: namely, relative to the propofol condition, the wake condition is comprised of epochs of linearly-predictable dynamics that were punctuated by sharp unpredictable deviations (Fig. 4f-g). These empirical and theoretical modelling results demonstrate that consciousness is concomitant with fluctuations in linear flow across the cerebral cortex, which are diminished by propofol anesthesia, however the question remains as to whether the thalamus itself plays a causal role in controlling these dynamics.

### Causal thalamic drive of cortical non-linear flow across conscious states

A decisive feature of biophysical models is that they can be used to create predictions that can be directly tested in empirical datasets across a range of imaging modalities and across species. We leveraged this feature to causally test the role of the thalamus in shaping the linearity of cortical dynamics. Specifically, we used an existing dataset comprising multi-electrode recordings from the frontal eye-fields and lateral interparietal cortical areas in a macaque monkey (Fig. 5b)^13,14^. Following propofol-mediated anesthesia, targeted 50Hz electrical stimulation of the matrix-rich central lateral thalamus drove the recovery of conscious arousal^13,14^. Despite the fact that the data was collected in a different species and through a different measurement modality that has a much faster temporal resolution than fMRI, we were able to use the generative nature of the biophysical model to create a testable prediction that mirrored the fMRI findings; Namely, that matrix stimulation should drive a recovery of the linearly unpredictable flow lost under propofol anesthesia (Fig. 5a) by flattening local attractors^11^. Importantly, this mechanism can be exploited while maintaining overall stability and facilitates long range interactions between specific brain areas^11,13^.

**Figure 5.**
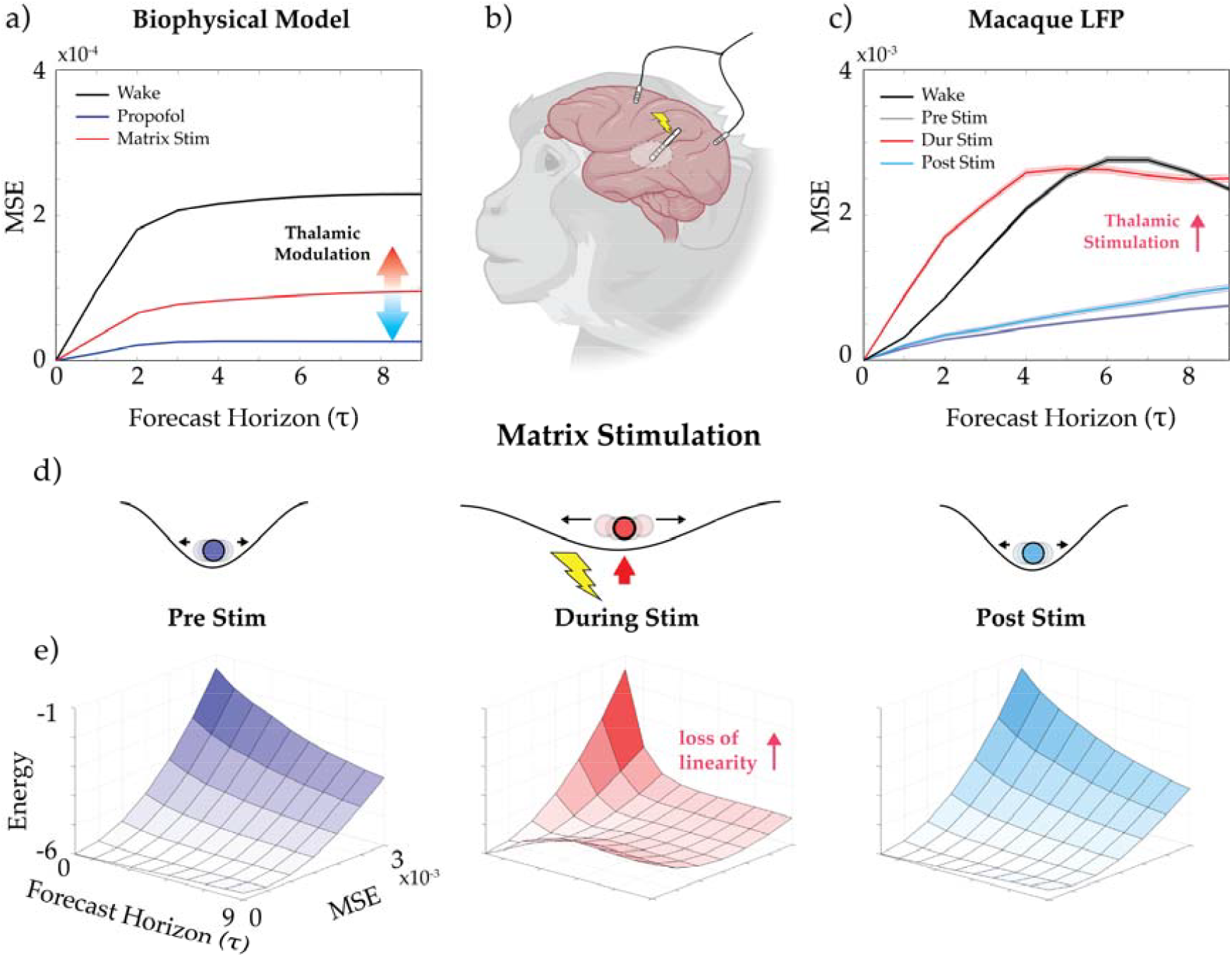
Causal thalamic influence on cortical linear dynamics. a) Using the corticothalamic biophysical model, we simulated matrix thalamic stimulation by applying a square-wave input to matrix-rich thalamic regions and then applied our linear flow analysis to the cortical electrophysiological time series during wake (black), following simulated propofol administration (blue) and following matrix stimulation (red) – matrix stimulation recovered non-linear cortical dynamics. b) experimental setup, in which a macaque monkey had 50Hz electrical stimulation applied to the matrix-rich central lateral thalamus while local field potentials were recorded from the frontal eye fields and lateral interparietal cortex. c) examination of the empirical recordings from macaque cortex recovered the predicted relationships: namely, central lateral thalamic stimulation recovered cortical non-linearity that matched the awake brain but was lost during propofol anesthesia. d) these effects are analogous to the global cortical state being enslaved to a deep attractor during propofol anesthesia (purple and blue) whose depth was diminished by matrix thalamic stimulation (red), allowing less linear cortical state dynamics. e) energy landscapes^62^ (i.e., the logarithm of inverse state probability) during pre-stim (purple), stimulation (red) and post-stimulation (blue) of the central lateral thalamus.

Using multi-electrode recordings from the macaque cortex, we confirmed that propofol had the predicted impact on cortical electrophysiology: namely, the induction of propofol increased linear predictability in the anesthetized monkeys, relative to wake (Fig. 5c), albeit on a shorter time-scale than the one observed in the fMRI data (though see Fig. S7). Next, we calculated this same linear predictability before (pre), during, and after (post) electrical stimulation of the central thalamus (which was effective in recovering the monkey’s conscious state; Fig. 5d). As predicted by our biophysical model, we observed substantial propofol-mediated linearity in the pre-stimulation window, a significant decrease in linear dynamics during stimulation (comparable to that seen in the wake condition), and a subsequent return to relatively predictable, linear dynamics following stimulus cessation (Fig. 5d). In addition, while the empirical stimulation recovered proportionally larger amounts of linear errors than the biophysical model (contrast the amplitude of the red lines in Fig. 5a and 5c), we anticipate that this difference is related to the simplified nature of the model, relative the nuanced neurobiological complexity of the brain. Irrespective of these details, our crucial prediction was confirmed: namely, that matrix thalamic stimulation reinvigorated deviations from linear cortical dynamics lost during propofol anesthesia.

As a final analysis, we again utilized the likelihood of changes in linear dynamics across each stimulation condition to formulate a statistical energy landscape (akin to the results observed in Fig. 3f-h). Analogous to an attractor landscape, this approach highlights the stability of on-going predictable cortical linear flow in the pre- and post-stimulation conditions, which is then destabilized by matrix thalamic stimulation (Fig. 5d). Our findings demonstrate that the matrix thalamus drives linearly unpredictable cortical flow by destabilizing local attractors across different states of arousal and provides a key feature of flexible dynamics indicative of versatile and adaptive cognition.

## Discussion

Here we showed that the thalamus controls the manner in which cortical dynamics fluctuate between laminar and non-laminar flow across states of consciousness. Using multimodal empirical neural recordings, a sensitive method from fluid dynamics and a biophysical model of the thalamocortical system, we demonstrated how a key principle of cellular organization within the thalamus controls these on-going linear dynamics, and specifically how the diffusely-projecting matrix thalamus destabilizes linear cortical flow. This approach reveals a candidate neuroanatomical mechanism for facilitating the adaptive cognitive benefits of the waking brain: namely, by facilitating brain dynamics that can support the formation of both stable and unstable brain states, which in turn can support reconfiguration as contingencies change across different behavioural contexts.

Previous work has shown how electrical stimulation of the thalamus can modulate conscious arousal^14,15^ and implicated the matrix thalamus as driving quasi-critical brain state formation^11,13^, thereby facilitating the recovery of conscious awareness from anesthesia. Our present findings enrich these insights and demonstrate how the matrix (diffuse)-core (targeted) cortical projections play a key role in shifting the cortex between laminar and non-laminar dynamics. Our results also represent a key capacity inherent to biophysical models, which is the ability to create mechanistic simulations that allow translations between different imaging modalities and recordings from different species. We anticipate that the addition of further anatomical constraints will only enrich the capacity for these models to provide additional testable predictions, though care must be taken to match the emergent dynamics of these models with the idiosyncratic spatiotemporal constraints imposed by standard cognitive contexts in which higher-order brain functions are interrogated.

Human neuroimaging studies have revealed a hierarchy of temporal and spatial autocorrelation scales across the cortex^49^ and that these two features capture a large number of existing topological metrics^22^. Our present findings expand these insights by quantifying intrinsic dynamic modal timescales that drive linear flow in neural activity across imaging modalities. The resulting spatiotemporal dynamic modes are defined by a complex (i.e., imaginary) eigenvector and corresponding eigenvalue^32,35,50^. Each mode contains four key characteristic measures: the spatial eigenvector defines a spatial pattern of coherent activity, and a corresponding spatial pattern of delays relative to each of these coherence patterns – called a dephasing map. In addition, each mode has a corresponding eigenvalue defining its oscillatory frequency, and an exponential gain parameter – which determines whether a mode will grow or decay in time (i.e., captures its temporal stability). The investigation of these features provides rich information regarding the types of dynamic modes that characterize the system. Our results also confirm the long-held notion that BOLD dynamics are predominantly constituted by slow and sustained patterns of activity (hence the popularity of the standard resting state network analyses^74,75^), but extend our ability to characterize and explore the spatial and temporal properties of the modes shaping these dynamics. We also demonstrate that the same anatomical mechanism underpins similar observations in both BOLD and electrophysiological data, which thus paves the way forward for future mechanistic work. Furthermore, our comparison of empirical imaging data to surrogates conserving covariance and Fourier spectra shows these features are not sufficient to explain the observed linear dynamics. An important open question that we were unable to interrogate in these data is whether non-neural sources of noise, such as physiological activity or task contexts, alters the observed relationship between thalamic activity and cortical dynamics, however this line of questioning is left for future, targeted experimental approaches.

It is worth noting that in this implementation, a linear model is generated for each subject over the complete length of that recording. As with calculation of covariance and Fourier power spectrum, this may also be performed at a finer temporal granularity via windowing approaches – this will favour faster timescale effects over slower ones, and thus should depend on the investigators question of interest. Here, the imaging data has been collected during the resting-state, but recordings with specific epochs of interest, such as during a task paradigm, would be served better by a piece-wise approach defining the linear dynamics instantiated by the brain within each epoch, particularly given the time-scale constraints imposed by the vast majority of task-based neuroimaging analysis protocols. Furthermore, our approach allows for the quantification of linear dynamics within a given timeseries, however, it does not make explicit claims about the nature of the residual dynamics. This is because these residuals may be multifaceted. Indeed, it may be that deviations from linearity are driven by non-linearities (in the mathematical sense) within the dynamical system becoming significant, due to the presence of extrinsic or intrinsic drive (i.e., visual/auditory/olfactory stimuli in the brain), or through the instantiation of a different transient linear system on a shorter timescale relative to the epoch of time initially considered. These are both exciting opportunities to be explored in future work.

In conclusion, we have demonstrated fluctuations in globally linear cortical dynamics in multimodal neuroimaging and electrophysiological data across human and non-human primates differentiates conscious arousal states and are controlled by the organization of the thalamus. In this way, we argue that this key feature of the brain’s neurobiological architecture helps to shape the robust, yet flexible, adaptive neural dynamics required for effective cognitive function that define our waking lives.

## Acknowledgements

E.J.M. was supported by an ARC DECRA Fellowship.

J.M.S. was supported by an NHMRC Investigator Fellowship.

## Methods

### Human neuroimaging data

Sixty healthy adult participants (28 females; 18–33 years; right-handed) were recruited and the research was approved by The University of Queensland Human Research Ethics Committee. These data were originally described in ^43^. 1050 (~10 minutes) whole-brain 7T resting state fMRI echo planar images were acquired using a multiband sequence (acceleration factor = 5; 2 mm^3^ voxels; 586 ms TR; 23 ms TE; 40^°^ flip angle; 208 mm FOV; 55 slices). Structural images were also collected to assist functional data pre-processing (MP2RAGE sequence – 0.75 mm^3^ voxels 4,300 ms TR; 3.44 ms TE; 256 slices).

DICOM images were first converted to NIfTI format and realigned. T1 images were reoriented, skull-stripped (FSL BET), and co-registered to the NIfTI functional images using statistical parametric mapping functions. Segmentation and the DARTEL algorithm were used to improve the estimation of non-neural signal in subject space and the spatial normalization. From each grey-matter voxel, the following signals were regressed: linear trends, signals from the six head-motion parameters (three translation, three rotation) and their temporal derivatives, white matter, and CSF (estimated from single-subject masks of white matter and CSF). The aCompCor method (Behzadi et al., 2007) was used to regress out residual signal unrelated to neural activity (i.e., five principal components derived from noise regions-of-interest in which the time series data were unlikely to be modulated by neural activity). Participants with head displacement > 3 mm in > 5% of volumes in any one scan were excluded (*n* = 5). A temporal band pass (0.001 < *f* < 0.125 Hz) was applied to the data. Following pre-processing, the mean time series was extracted from 400 pre-defined cortical parcels using the Schaefer atlas (Schaefer et al., 2018).

For the anesthesia BOLD fMRI dataset, we utilized data derived from previously published research works^52,76^ that have been openly shared on the OpenNeuro data repository (doi:10.18112/openneuro.ds003171.v2.0.0). The dataset comprises 17 healthy individuals (Age: 24± 5, M/F: 13/4), all of whom were right-handed, native English speakers, and had no recorded history of neurological disorders. The original study obtained ethical approval from both the Health Sciences Research Ethics Board and Psychology Research Ethics Board of Western University (REB #104,755) and adhered to the principles outlined in the revised declaration of Helsinki (2000). We analyzed two conditions, awake (fully alert and communicative) and deep sedation, which was achieved with an initial target effect-site concentration of 0.6 µg/ml and oxygen titrated to maintain SpO2 above 96%, with increments of 0.3 µg/ml with repeated assessments of responsiveness until deep sedation (Ramsey level 5).

Imaging was performed on a 3T Siemens Tim Trio system with a 32-channel head coil. Subjects resting state fMRI scans using BOLD EPI sequence (33 slices, voxel size: 3mm^3^ isotropic, inter-slice gap of 25%, TR = 2000 ms, TE = 30 ms, matrix size = 64×64, FA = 75°). Resting-state scans had 256 volumes. Anatomical scans were also obtained using a T1-weighted 3D Magnetization Prepared - Rapid Gradient Echo (MPRAGE) sequence (voxel size: 1mm^3^ isotropic, TR = 2.3, TE = 4.25 ms, matrix size = 240 × 256 × 192, FA = 9°).

Preprocessing was completed using *fMRIPrep* standard pipeline which involves the basic preprocessing steps (co-registration, normalization, unwarping, noise component extraction, segmentation, skull stripping, etc.). The extracted time series were denoised specifying motion and physiological signals from white matter and CSF, with high-pass and low-pass band filters set at 0.01 and 0.1, respectively. The global signal was removed by standardizing each time point^39^.

Thalamic timeseries representing the ‘core’ and ‘matrix’ populations were estimated by estimating the dot-product between the voxel-wise time-series of thalamic BOLD signal and a standardized map of the relative mRNA expression levels for PVALB (‘core’) and CALB1 (‘matrix’) provided by the Allen Human Brain Atlas^8,9,77^. Variogram modelling was applied prior to our obtaining the gene maps to correct for potential subject-level differences^78^. Note that there are other calcium binding proteins with non-trivial expression in the thalamus^61,79^, however these patterns were not considered here, as they have not been directly linked to the same core-matrix gradient in the thalamus^9^. Note that we do not advocate for a strict dichotomy between core and matrix neurons in this work, but rather that these two projection types represent anatomical extremes at either end of an approximately continuous spectrum^54,56^. See Muller et al 2020^8^ for more details.

### Macaque electrophysiological dataset

We compared modelling data to electrophysiology data from our previous study^14^. This dataset consists of simultaneous local field potential (LFP) recordings from the frontal eye field (FEF), lateral intraparietal area (LIP) and central lateral thalamus (CL) in the right hemisphere of two macaques (*Macaca mulatta*, 4.3-5.5 years old, 7.63-10.30 kg body weight). We lowpass filtered LFPs to 250 Hz, then linearly detrended and extracted artifacts. Bipolar derivations of the LFPs were then calculated to minimize any possible effects of a common reference and volume conduction. Recordings were performed during general anesthesia – either isoflurane (0.8%–1.5% on 1 L/min O2 flow; 9 sessions) or propofol (0.17-0.33 mg/kg/min i.v.; 9 sessions) – or wakefulness, using 16- or 24-contact linear micro-electrode arrays (LMEAs; MicroProbes). The electrode contacts had a diameter of 12.5µm, and 200µm spacing between contacts.

We performed deep brain stimulation via simultaneous stimulation of 16 contacts of the LMEAs in the central thalamus. We applied 400 ms bi-phasic pulses of 200 mA, at 50 Hz stimulation frequency, for a total of 60 s stimulation duration for any given stimulation event. To localize LMEAs, we averaged two 3D T1-weighted structural images of the MRI-compatible electrodes *in situ* (inversion-recovery prepared gradient echo sequence with: FOV = 128*mm*^2^; matrix = 256 × 256; no. of slices = 166; 0.5 mm isotropic; TR = 9.68 ms; TE = 4.192 ms; flip angle = 12°; inversion time (TI) = 450 ms) and cross-validated the electrode location with electrode depth measurements during recordings/stimulation. We used an arousal index (0-10) based on eye openings, body movements and vital signs, as well as EEG and EMG, to measured stimulation-induced changes in the level of consciousness. We defined effective stimulations, increasing the level of consciousness, as having an arousal index of ≥3, which all experimenters could differentiate from ineffective stimulations having an arousal index of 0-2. See ^14,80–82^ for more details. The University of Wisconsin – Madison Institutional Animal Care and Use Committee approved all procedures, which conformed to the National Institutes of Health Guide for the Care and Use of Laboratory Animals. Due to hardware constraints, the LFP time-series were down-sampled from 1kHz to 100Hz for linear model estimation – although we note that results in Fig. 5c and subsequent conclusions are not timescale sensitive as shown in Supp. Fig. 7.

### Linear Forecast Analysis (LFA)

Here we outline LFA as applied to all datasets within this manuscript. First, each timeseries Z= [**z**_1_ **z**_2_ **z**_3_ … **z**_*n*_] with *n* observations of *p* variables is partitioned into a pre x and post Y timeseries – where:

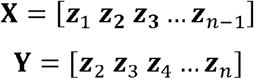

with x lacking the final time point of Z and Y lacking the initial time point. Note we have dropped the function-of-time notation for simplicity. Singular value decomposition is then applied, **x = U × s × v^T^**, and a rank reduction of the data is made that captures 95% of the explained variance. Although not explicitly necessary for timeseries with significantly more timepoints than variables, some of the data used here violate this condition and thus the results are standardized around this threshold – see Supp. Fig. 4 showing truncation effects on eigenspectrum. A linear propagator matrix **A** can be estimated from the data where **Y = Ax** as follows:

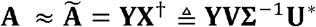

By stepping through each timepoint of **Z** projected into the SVD sub-space, the linear propagator can be used to forecast the proceeding time points. Mean-squared-error is used to compare between the predicted timeseries and the ground-truth timeseries:

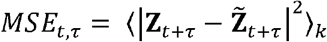

where Z is the ground-truth data and 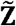 is the forecasted timeseries using the linear model, and *k* is the number of SVD modes.

### Dynamic mode decomposition

The eigen decomposition of the linear propagator **A** gives the dynamic modes of the system, **Aw = WΛ** where **w** is the matrix of eigenvectors, and **Λ** is the diagonal matrix of eigenvalues λ*i*. The DMD approximation of the data can then be written as the simple dynamic model:

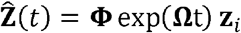

where **Ω ≜ YVΣ-1w** with **Ω** = log (Λ)/Δt and t is time^35^. **z**_*i*_ is the initialization state and can be solved for each timepoint using the pseudoinverse.

Surrogate models

The spectral surrogate utilized here is adapted from ^21^ and generates a white noise timeseries with the same dimensions as the empirical data. The Fourier transform is then applied to both the surrogate and empirical timeseries, and these are multiplied together in this spectral domain. Following the inverse Fourier transform, the resulting timeseries is projected through the eigenvectors of the empirical covariance matrix to give a null model time series with matched covariance and Fourier power spectrum.

The shuffled surrogate is generated by performing a uniform scrambling of the empirical time-series indices. This ensures regional co-activations at each time point are conserved, but their temporal position relative to activations in scramble – i.e., the same covariance matrix but scrambled moment-to-moment changes.

### Linear mode clusters

k-means clustering is used to define clusters of linear mode timescales and spatial coherence independently. In both cases, a sweep of cluster sizes k=2-20 with 100 repetitions for each is performed. A peak in adjusted mutual information (calculated using BCT http://www.brain-connectivity-toolbox.net/) between each repetition’s clusters for the same cluster size is used to select a cluster size of k=5. Having selected this cluster size a final run of 1000 repetitions is used to find the optimal clustering. Timescale clustering is performed on the eigenvalues of all subjects (N=59) linear modes, i.e., the 2-dimensional complex numbers defining damping rates and oscillation frequencies. Spatial clustering is performed on the coherence maps defines by each linear mode across all subjects (N=59), i.e., the real component of the complex eigenvectors of the linear propagator.

### Energy landscapes

Following methodology from previous work^62^, we formulate an energy landscape by calculating the probability of observing a given linear model error (MSE) across the entire timeseries using a Gaussian kernel density estimation,

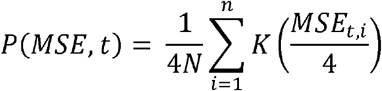

where 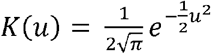. As is typical in statistical mechanics the energy of a given state, Eσ, and its probability are related by 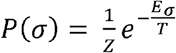 where *Z* is the normalization function and *T* is the scaling factor equivalent to temperature in thermodynamics^62^. In our analysis Σσ Pσ= 1→Z = 1 by construction and we can set *T*= 1 for the observed data. Thus, the energy of each MSD at a given time-lag,, is then equal to the natural logarithm of the inverse probability, *P*(*MSE,t*) of its occurrence,

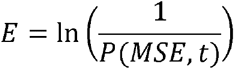

### Corticothalamic model

The corticothalamic model consists of 400 coupled neural masses. We outline this architecture by first detailing the corticothalamic neural mass as follows. The corticothalamic neural mass model used in this work contains four distinct populations: an excitatory pyramidal cell, *e*, and an inhibitory interneuron, *i*, population in the cortex; and two excitatory nuclei, matrix, *s*_*m*_; core, *s*_c,_ and inhibitory thalamic reticular nuclei, *r*, population in the thalamus (Fig. 1c). The dynamical processes that occur within and between populations in a neural field model are defined as follows:

For each population, the mean soma potential results from incoming postsynaptic potentials (PSPs):

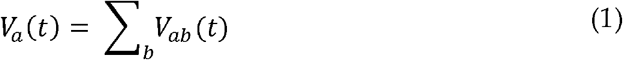

where *V*_*ab*_ (*t*) is the result of a postsynaptic potential of type *b* onto a neuron of type *a* and *a,b* ∈ {*e,i,r,s*}. The postsynaptic potential response in the dendrite is given by

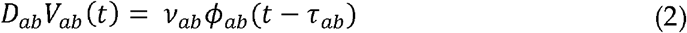

where the influence of incoming spikes to population *a* from population *b* is weighted by a connection strength parameter 𝓋_*ab*_ = *N*_*ab*_*S*_*ab*_, with the mean number of connections between the two populations *N*_*ab*_ and *s*_*ab*_ is the mean strength of response in neuron *a* to a single spike from neuron *b*. τ_*ab*_ is the average axonal delay for the transmission of signals, and ϕ_*ab*_ is the mean axonal pulse rate from *b* to *a*.

The operator *D*_*ab*_ describes the time evolution of *V*_*ab*_ in response to synaptic input,

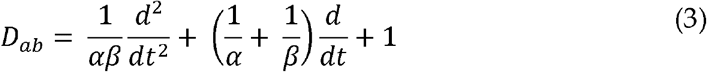

where *β* and *α* are the overall rise and decay response rates to the synaptodendritic and soma dynamics.

The mean firing rate of a neural population *Qa*(*t*) can be approximately related to its mean membrane potential, *Va* (*t*), by

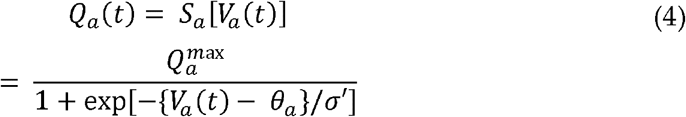

which define a sigmoidal mapping function *S*_*a*_ a with a maximal firing rate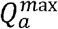, a mean firing threshold θ_*a*_, and a standard deviation of this threshold 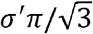.

The mean axonal pulse rate is related to the mean firing rate by,

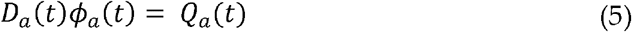

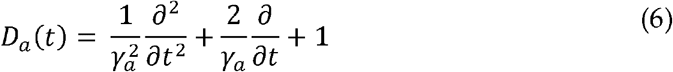

Here, γ*a* = 𝓋_*a*_/*r*_*a*_ represents the damping rate, where 𝓋_*a*_ is the propagation velocity in axons, and *ra* is the characteristic axonal length for the population.

A network of 400 corticothalamic neural masses were simulated using the neural field simulation software, *NFTsim*^*83*^. The parameters for each neural mass were identically set to *“eyes-closed”* estimates given in Table 1^68,69,71,83,84^, which results in simulated activity with a 1/f spectrum and a peak in the alpha frequency band (8-13 Hz) in the absence of network coupling. These are example parameters representative of the “*eyes-closed”* state following Bayesian model fits to human EEG power spectra^69^. Many preceding studies^68,69,71,72,84–86^ have shown the linear transfer function, which drives the linear spectral content of the corticothalamic model, is derived and shown to be low-dimensional, i.e., only a few loop gains in the system are needed to capture the key features of the power spectra. In this particular context, the eyes-closed state of human EEG has a 1/f slope and a spectral peak in the 8-13Hz alpha frequency band, and this is explained by a weakly damped thalamocortical loop gain (ese). For the present study we have selected characteristic parameters for this power spectrum (Table 7 from ^69^). Note that this spectrum describes modulations of firing rate around a fixed point, which are static for the network. These spike rate modulations in neurons drive changes in the extracellular electric field which are then measured via EEG and LFP recordings^87^. In this way, we are able to compare our model outputs to empirical data through the power spectral density function.

**Table 1.** Corticothalamic neural mass parameters. Adapted from ^69^.

| ***Parameter*** | ***Description*** | ***Value*** | ***Unit*** |
| --- | --- | --- | --- |
| $\gamma_e$ | Cortical damping rate | 116 | s<sup>-1</sup> |
| $Q^{max}$ | Maximum firing rate | 340 | s<sup>-1</sup> |
| $\theta$ | Firing threshold | 12.9 | mV |
| $\sigma'$ | Threshold spread | 3.8 | mV |
| $\phi_n$ | Input noise amplitude spectral density | $1 \times 10^{-5}$ | s<sup>-1</sup> |
| $\alpha$ | Decay rate of cell-body potential | 83 | s<sup>-1</sup> |
| $\beta$ | Rise rate of cell-body potential | 769 | $s^{-1}$ |
|  | Intra-node coupling strengths |  |  |
| $\nu_{ee}$ | | 1.5 | mV s |
| $\nu_{ei}$ | | -3 | mV s |
| $\nu_{esc}$ | | 0.57 | mV s |
| $\nu_{se}$ | | 3.4 | mV s |
| $\nu_{scr}$ | | -1.5 | mV s |
| $\nu_{scn}$ | | 3.6 | mV s |
| $\nu_{re}$ | | 0.17 | mV s |
| $\nu_{rsc}$ | | 0.05 | mV s |
| $\tau_{esc,esm} + \tau_{sce,sme}$ | Corticothalamic loop delay | 85 | ms |

Each simulation was run for a total of 64s with 7.5s of initial transients removed using an integration timestep of Δ*t* = 2-l3s. This minimizes contributions from the model’s initial state and ensures the integration algorithm has stabilized before we begin analysing simulation outputs. Longer simulations produced qualitatively identical results, as did shorter simulations, however, many of the analysis measures presented in this paper perform better with more data – i.e., correlation and coherence are noisier with less data. Thus, a balance between metric accuracy, resource allotment, and tractability for dataset manipulations was chosen. All outputs were down-sampled to 200Hz for tractability. All remaining data were used for subsequent analysis.

### Structural Connectivity

The structural connectivity used to define the model network consists of a combination of distance dependence and long-range connectivity estimated from white-matter fibre densities measurement^88^. The distance dependence was generated via an exponentially decreasing function. First, the geodesic distance between all nodes, which correspond to parcels from^89^ with MNI coordinates, are calculated along the *fsaverage* cortical surface mesh^90^ using the Fast-Marching algorithm (Gabriel Peyre (2022)ToolboxFastMarching:https://www.mathworks.com/matlabcentral/fileexchange/6110-toolbox-fast-marching). The geodesic distances are then scaled as an exponentially decreasing function of distance,

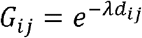

where *d*_*ij*_ is the geodesic distance in MNI space along the surface mesh and λ is the decay rate. A 200×200 connection matrix is defined in this way for each hemisphere. We further assume that interhemispheric connectivity is symmetric and one-to-one, and thus the full 400×400 distance dependence network is composed as

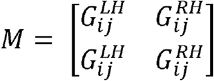

Where 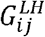 and 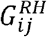are the left and right interhemispheric connectivities, respectively.

The complete network connectivity is then formulated via the summation of the distance dependence matrix and the empirically estimated white-matter connectivity (both normalized by their respective maximum values). Since the strength of these connections is not known empirically, we follow other approaches and sweep values of a global scaling of this hybrid connection matrix, as well as the proportion of distance dependence-to-white-matter connectivity and distance decay rate parameter. Functional connectivity of these parameter sweeps is then compared to empirical resting-state BOLD data ^91^ to define optimal values (see Supp. Fig. 2)

### Model balancing

In order to maintain stability in the model, excitatory inputs to a given node, coupled via the structural network connections, must be balanced with a corresponding inhibition. To do this, we first compute the total excitatory connection strength to each corticothalamic node.

We then leverage an assumption from previous neural field models, namely that excitatory and inhibitory synapses in the cortex can be assumed proportional to the number of neurons ^*86,92*^. This random connectivity approximation results in 𝓋_ee_ = 𝓋_*ie*_, and 𝓋_*ei*_ = 𝓋_*ii*_ which implies *V*_*e*_ = *V*_*i*_ and *Q*_*e*_ = *Q*_*i*_. Inhibitory population variables can then be expressed in terms of excitatory quantities. Whilst we do not make this assumption in the present model, we can leverage it to refine an inhibition scaling that balances the excitatory inputs from our specific structural network.

In the reduced corticothalamic neural mass, the fixed-point attractors, or steady states are found by setting all time derivatives in the above equations to zero. The steady-state values 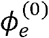of ϕ_*e*_ are then given by solutions of

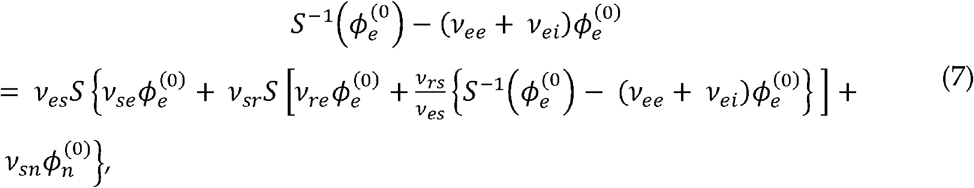

where 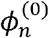 is the steady state component of the input stimulus ^*92,93*^. Roots of Eq. (7) are found using the *fzero()* function from MATLAB.

Following similar approaches^94^, we leverage Eqn. 7 by setting 𝓋_*ee*_ equal to each corticothalamic neural masses network coupling defined by the structural connectivity. We then set the cortical firing rate to be 3Hz, in-line with empirical observations, and numerically solve for cortical inhibition 𝓋_*ei*_. This results in each neural mass having a 3Hz steady-state cortical firing rate across the network, despite having heterogeneous network connectivity.

Note that the diffuse matrix inputs to each corticothalamic neural mass are excluded from this balancing as they are purely excitatory and only target the excitatory cortical population. Thus, the overall effect of matrix inputs is to distort each local attractor, increasing their firing rates when they are coupling to the network^11^. In addition, matrix nuclei are known to project to the reticular thalamic nucleus in rodents ^9,95–97^, albeit weakly^98^, but have been excluded from the current model for simplicity.

### Modelling Propofol

The effect of propofol is modelled as an up-regulation of GABA-a receptors which prolongs inhibitory postsynaptic response potentials. This is implemented as an increase to the synaptodendritic functions (Eqn. 3) decay rate parameter, α, for all inhibitory connections in the corticothalamic neural mass. In addition, consistent with previous approaches^99^, we maintain a constant peak amplitude of the IPSP functions following the rescaling. The solution to Eqn. 3 for a delta function input corresponds to,

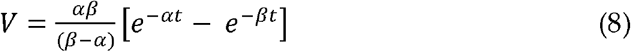

which has a peak amplitude at 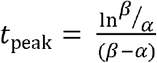. The rescaling of 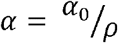 defines a new peak potential at *t*_peak_ which is renormalized to its pre-propofol value. We leverage the change in coherent alpha-band activity (8-13Hz) observed in ^14^ between LIP and FEF to optimize propofols effect in the model, which results in ρ = 1.127.

**Supp. Figure 1.**
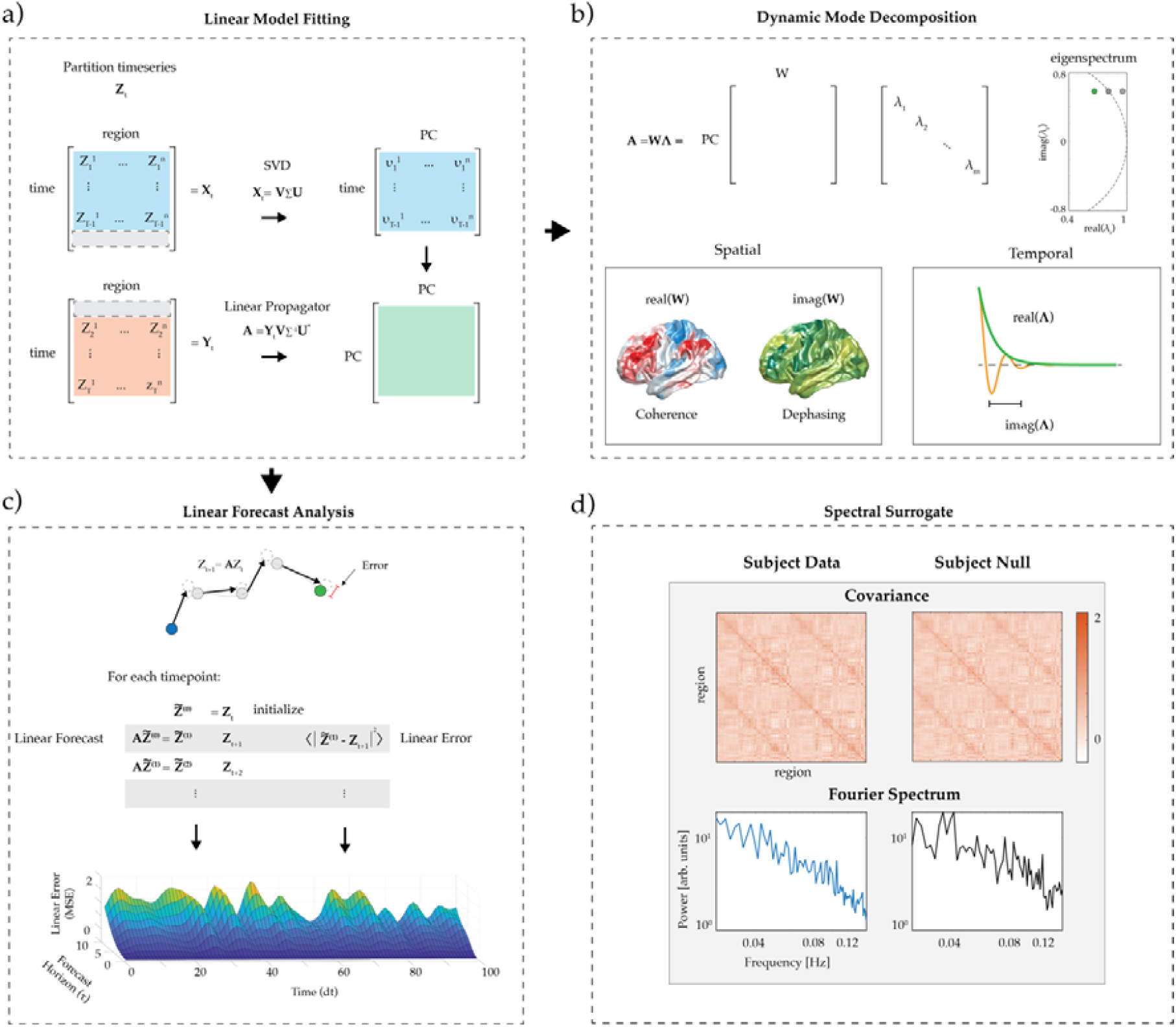
Linear Forecasting and Spectral Surrogate. a) Formulation of Linear Propagator via the singular-value decomposition. b) Dynamic mode decomposition – eigenmode decomposition of linear propagator matrix resulting in oscillatory spatiotemporal modes. c) Forecast Horizon Analysis using the current state and the linear propagator matrix to predict future time points. Mean-squared-error is used to compare forecast dynamics to ground-truth empirical timeseries. d) Example spectral surrogate for a single subject showing matched covariance and Fourier power spectrum.

**Supp. Figure 2.**
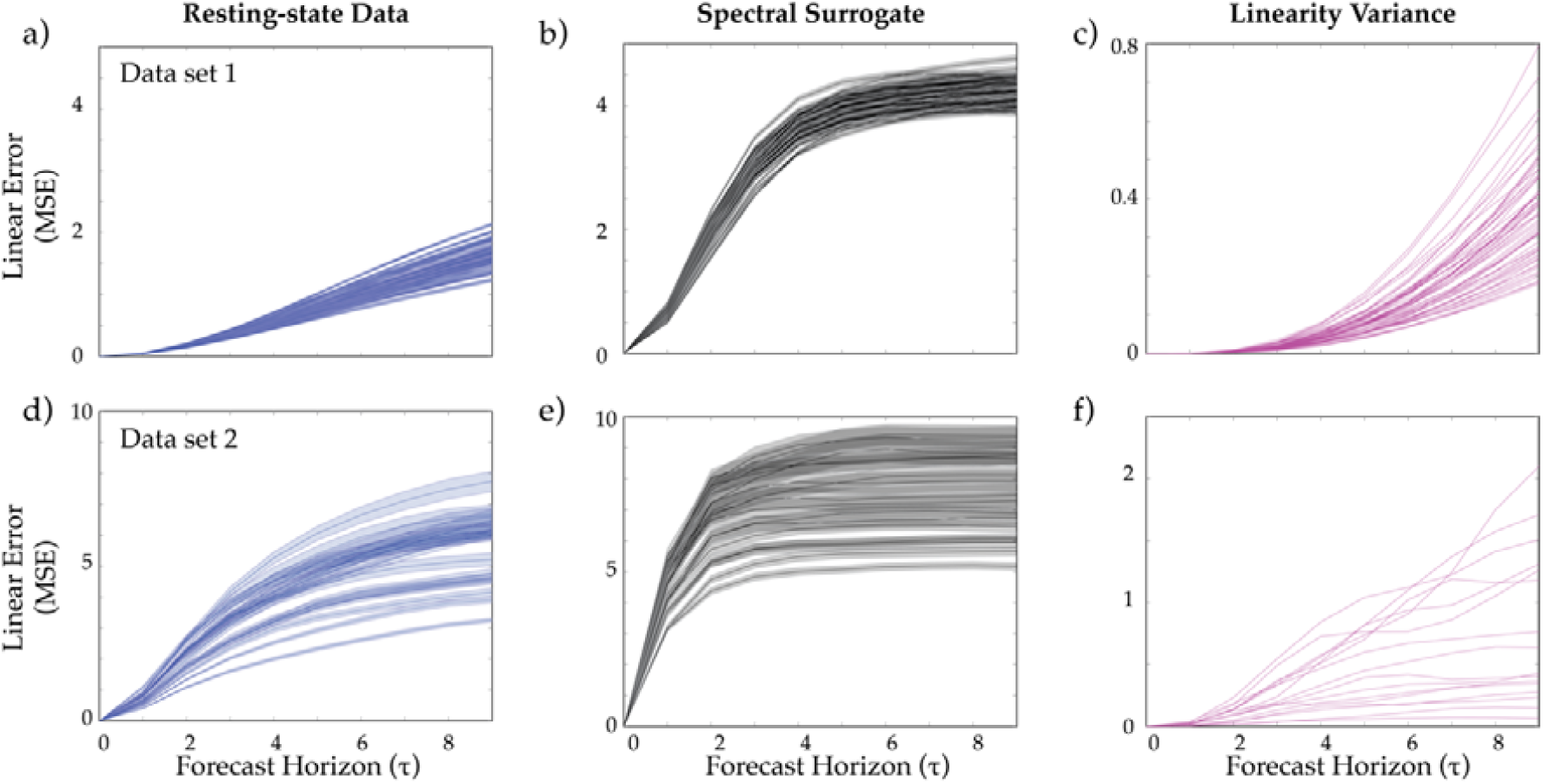
LFA cross-validation. First Row: 7T human resting-state BOLD data^43^ (N=59). a) Linear Error for each forecast horizon of the empirical data. b) Spectral surrogate for each subject. c) Variance of linear error (MSE) and a function of forecast horizon. Second Row: 3T human resting-state BOLD data^52^ (N=14) d) Linear Error for each forecast horizon of the empirical data. e) Spectral surrogate for each subject. f) Variance of linear error (MSE) and a function of forecast horizon.

**Supp. Figure 3.**
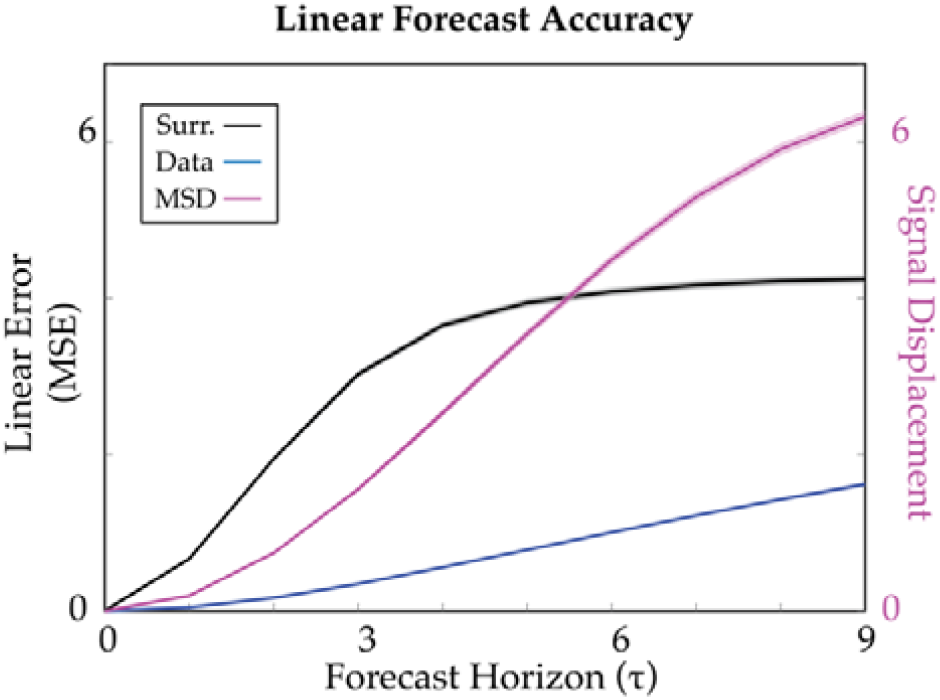
Linear Model Performance. Left axis: Linear error as a function of forecast horizon for the subject average 7T resting-state human imaging data(blue), spectral surrogate(black). Right axis: Mean-squared displacement of the subject average 7T resting-state human imaging data showing general variability within the recordings.

**Supp. Figure 4.**
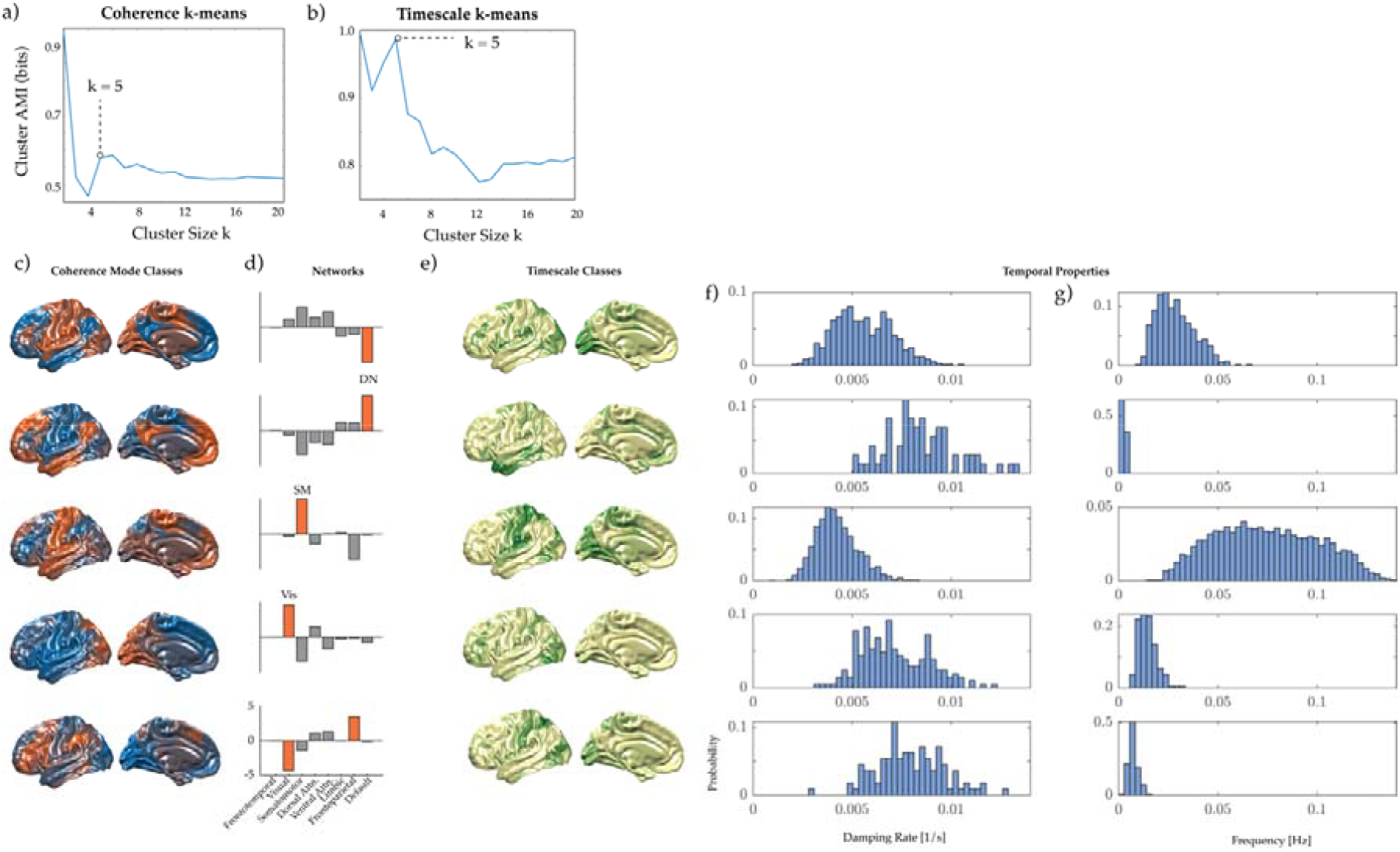
K-means optimization and Dynamic mode clusters. a) Coherence map clustering: Adjusted mutual information for 100 k-means repetitions for each cluster size k. b) Eigenmode clustering: Adjusted mutual information for 100 k-means repetitions for each cluster size k. c) Coherence cluster centroids d) Coherence cluster dot-product with resting-state networks e) Average coherence maps of eigenvalue clusters f) Timescale cluster probability distribution for damping rates. g) Timescale cluster probability distribution for oscillatory frequencies.

**Supp. Figure 5.**
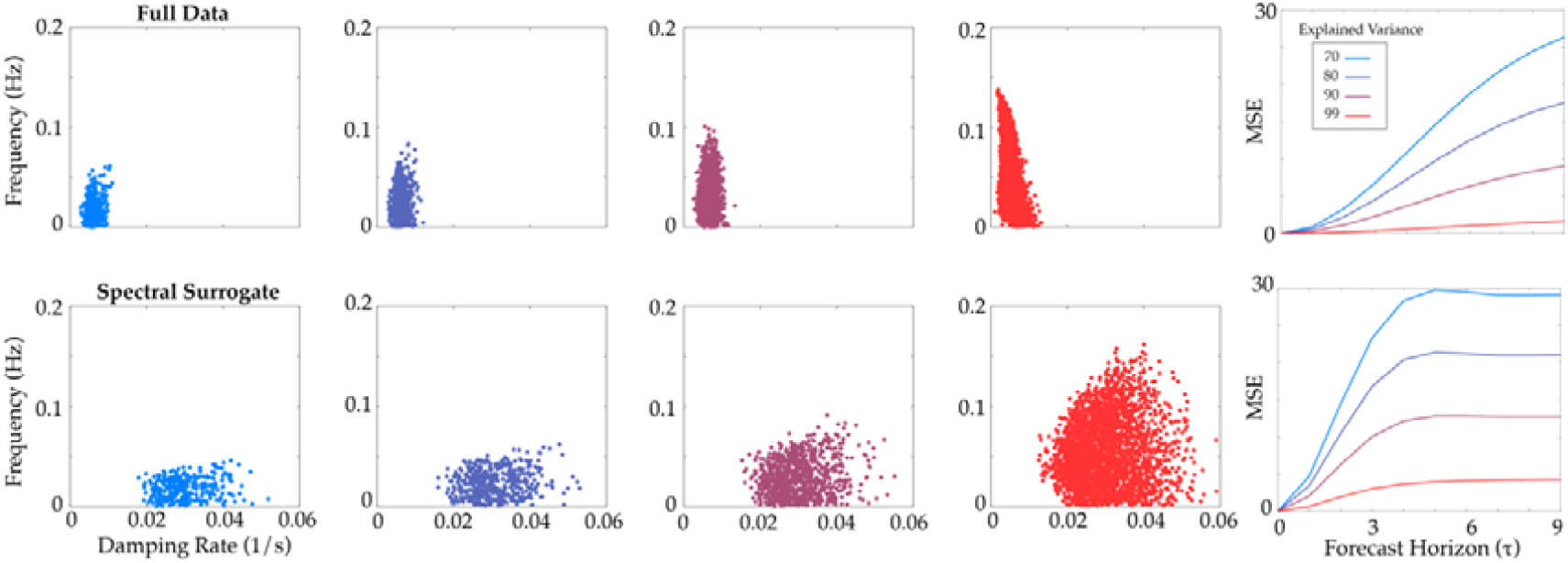
Linearity, dynamics modes and explained variance. Dynamic mode timescales and linearity for the a) full data (7T resting-state N = 59) and b) covariance and Fourier spectrum matched surrogates, as a function of the included variance explained.

**Supp. Figure 6.**
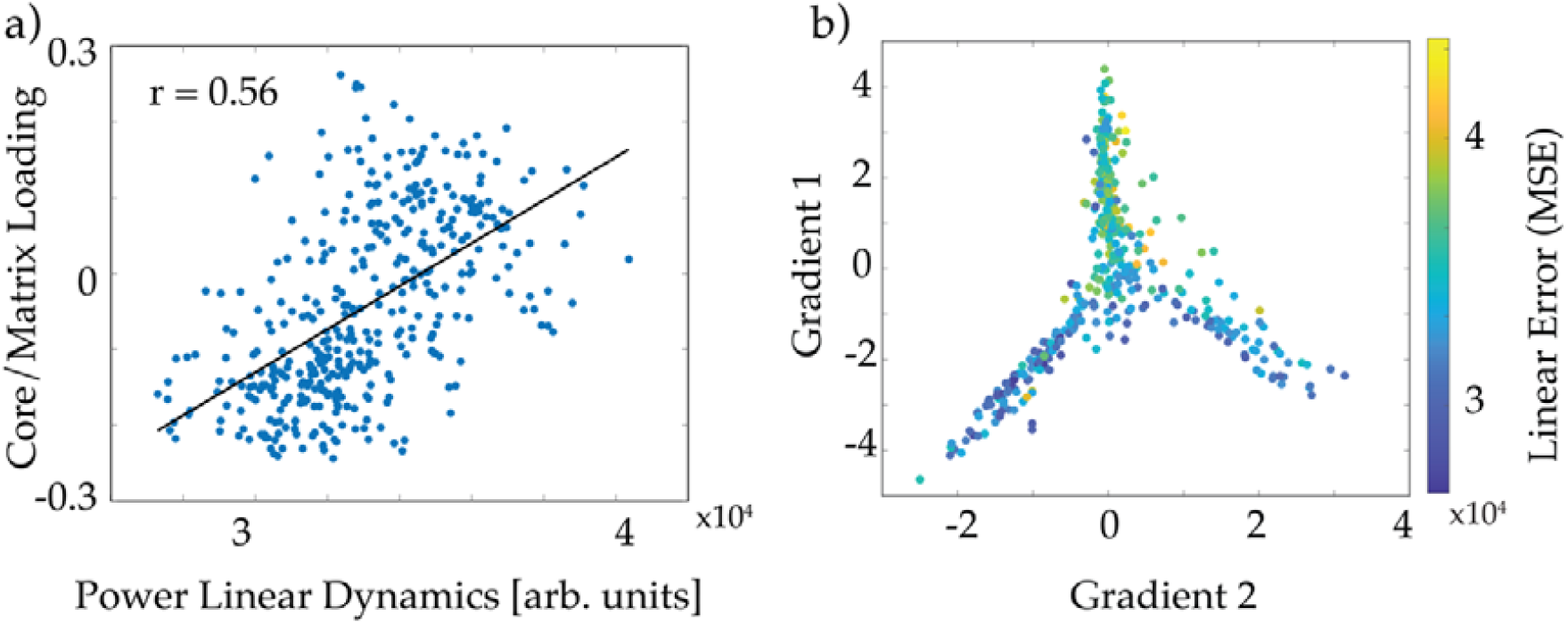
Linear cortical dynamics. a) Corticothalamic Core-Matrix correlation to linear cortical dynamics. b) Spatial loading of linear cortical dynamics to a low-dimensional gradient from_100_.

**Supp. Figure 7.**
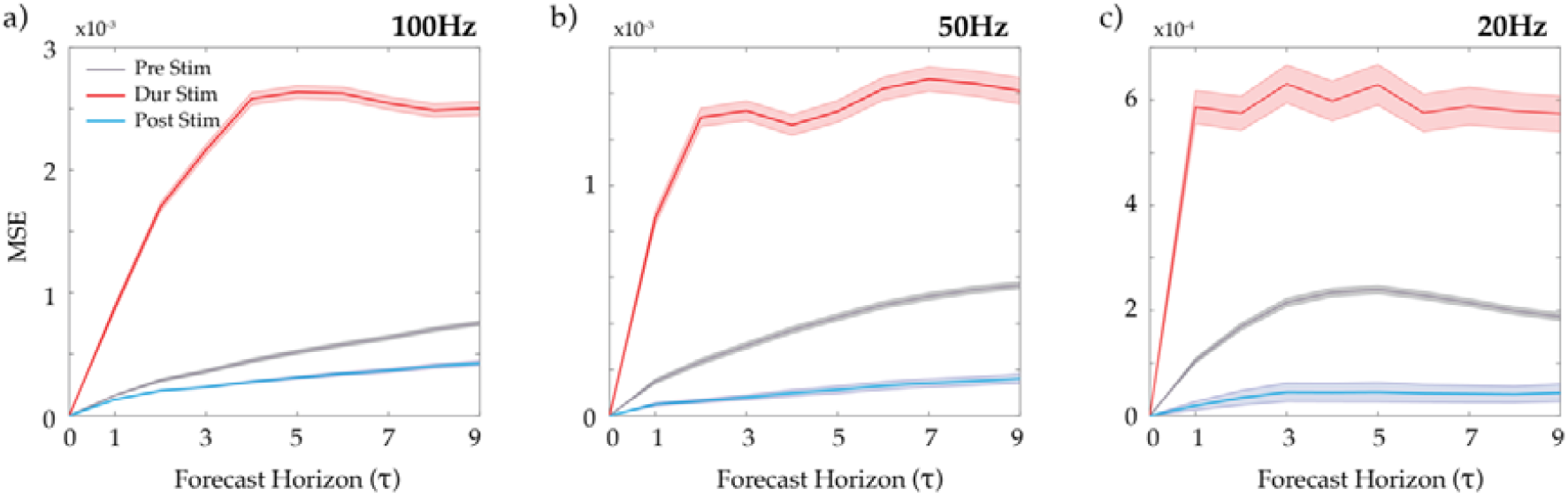
Monkey down-sampling effects. Linearity shifts of macaque multielectrode electrophysiological data pre, during, and post thalamic stimulation for a) 100Hz, b) 50Hz, and c) 20Hz sampling.

### Adaptive timescales and dynamic brain modes

Intrinsic timescales of the brain have been of interest since the discovery of oscillatory dynamics in the first human EEG recordings^48^. Human neuroimaging studies have revealed a hierarchy of temporal and spatial autocorrelation scales across the cortex^49^ and that these capture a large number of existing topological metrics^22^. Linear flow analysis enriches this approach by quantifying intrinsic modal timescales that drive linear flow in neural activity. The resulting dynamic modes are defined by a complex (i.e., imaginary) eigenvector and corresponding eigenvalue^32,35,50^. Each mode contains four key characteristic properties: the spatial eigenvector defines a spatial pattern of coherent activity, and a corresponding spatial pattern of delays relative to each of these coherence patterns – called a dephasing map. In addition, each mode has a corresponding eigenvalue defining its oscillatory frequency, and an exponential gain parameter – which determines whether a mode will grow or decay in time (i.e., captures its temporal stability). The investigation of these features provides rich information regarding the types of dynamic modes that characterize the system.

In order to gain insight into how these dynamic modes relate across individual subject recordings, we leverage a two-pronged clustering approach to aggregate modes across timescales, and spatial coherence, respectively. First, we consider timescales by utilizing *k*-means clustering of the eigenspectrum collated across all subject recordings. A peak in adjusted mutual information between clustering repetitions was then used to select *k* = 5 clusters (1000 repetitions; see Supp. Fig. 4), resulting in 5 distinct timescale groupings – the corresponding average of the coherence maps within these clusters (Supp. Fig. 4). Timescale cluster 1 shows somatomotor and visual activation antiphase with frontal, parietal and temporal cortices across a broad range of frequencies – spanning much of the time resolution of BOLD recordings. These modes are also weakly damped – i.e., have a strong gain relative to the other temporal clusters. Timescale cluster 2 shows visual, premotor, and dorsal lateral prefrontal antiphase with other cortices at slower frequencies (< 0.03 Hz; Fig. g-h middle) and with a stronger damping rate, i.e., weaker gain, than cluster 1. And timescale cluster 3 shows strongly damped temporal cortex activity at frequencies < 0.02 Hz.

Next, we considered the spatial patterns of coherence, defined by each mode, aggregated across subjects and recording sessions to define consistent modal groupings^50^. Again, we leveraged *k*-means clustering (adjusted mutual information peaked between clustering repetitions for *k* = 5; See Supp. Fig. 3): Figure 2i shows 3 cluster centroids and their defining coherence patterns – i.e., the spatial patterns of the linear modes across all subjects fall within distinct classes. These classes show a significant overlap with resting-state networks used broadly in the literature^51^ – including the default mode, and somatomotor and visual networks. We further find that the damping rates and frequencies of the modes within each class have comparable probability distributions – demonstrating these spatial classes did not demonstrate unique linear timescales at this granularity in resting-state.

